# Cas12a/Cpf1 knock-in mice enable efficient multiplexed immune cell engineering

**DOI:** 10.1101/2023.03.14.532657

**Authors:** Matthew B. Dong, Kaiyuan Tang, Xiaoyu Zhou, Johanna Shen, Krista Chen, Hyunu R. Kim, Jerry Zhou, Hanbing Cao, Erica Vandenbulcke, Yueqi Zhang, Ryan D. Chow, Andrew Du, Kazushi Suzuki, Shao-Yu Fang, Medha Majety, Xiaoyun Dai, Sidi Chen

**Affiliations:** Department of Genetics, Yale University School of Medicine, New Haven, Connecticut, USA; System Biology Institute, Yale University, West Haven, Connecticut, USA; Center for Cancer Systems Biology, Yale University, West Haven, Connecticut, USA; M.D.-Ph.D. Program, Yale University, West Haven, Connecticut, USA; Immunobiology Program, Yale University, New Haven, Connecticut, USA; Department of Immunobiology, Yale University, New Haven, Connecticut, USA; Molecular Cell Biology, Genetics, and Development Program, Yale University, New Haven, CT, USA; Yale College, Yale University, New Haven, Connecticut, USA; Department of Neurosurgery, Yale University School of Medicine, New Haven, Connecticut, USA; Yale Comprehensive Cancer Center, Yale University School of Medicine, New Haven, Connecticut, USA; Yale Stem Cell Center, Yale University School of Medicine, New Haven, Connecticut, USA; Yale Center for Biomedical Data Science, Yale University School of Medicine, New Haven, Connecticut, USA

**Keywords:** Cas12a/Cpf1, knock-in mice, immune engineering, gene editing

## Abstract

Cas9 transgenic animals have drastically accelerated the discovery of novel immune modulators. But due to its inability to process its own CRISPR RNAs (crRNAs), simultaneous multiplexed gene perturbations using Cas9 remains limited, especially by pseudoviral vectors. Cas12a/Cpf1, however, can process concatenated crRNA arrays for this purpose. Here, we created conditional and constitutive LbCas12a knock-in transgenic mice. With these mice, we demonstrated efficient multiplexed gene editing and surface protein knockdown within individual primary immune cells. We showed genome editing across multiple types of primary immune cells including CD4 and CD8 T cells, B cells, and bone-marrow derived dendritic cells. These transgenic animals, along with the accompanying viral vectors, together provide a versatile toolkit for a broad range of *ex vivo* and *in vivo* gene editing applications, including fundamental immunological discovery and immune gene engineering.

## Introduction

Immunotherapy has greatly expanded the oncological armamentarium. These therapies range from innate immune modulators that activate antitumor adaptive immune responses to genetically engineered tumor-specific T cells (Dougan et al., 2019; Waldman et al., 2020). Many of these therapies converge on the activation of T cells that can specifically detect and eliminate tumor cells (Waldman *et al*., 2020). Although these therapies have met tremendous clinical success in a broad range of tumor types, many patients fail to mount durable and sustained antitumor responses, leading to increased interest amongst the biomedical research community towards identifying additional critical modulators of antitumor immunity.

CRISPR-based gene editing technologies have accelerated the pace at which novel genetic regulators are being discovered because they can be programmed to target specific DNA sequences (Anzalone et al., 2020; Hsu et al., 2014; Komor et al., 2016). The accuracy and efficiency of these technologies have improved with the advancements made in CRISPR RNA (crRNA) library construction, next generation sequencing, and widely available computational pipelines (Doench, 2018; Shalem et al., 2014; Shalem et al., 2015). Of the many CRISPR tools that depend on RNA-guided endonucleases (RGNs), Streptococcus pyogenes Cas9 (SpCas9) and its variations have been the most widely used. Transgenic expression of Cas9 and Cas9-variants in mice enable safe and highly efficient *in vivo* and *ex vivo* genomic manipulation of primary cells by streamlining the pseudoviral vectors down to just crRNAs and transgenic reporter genes (Chu et al., 2016; Gemberling et al., 2021; Hunt et al., 2021; Platt et al., 2014; Zhou et al., 2018). Collectively, RGN transgenic animals and the improvements in crRNA design have enabled pooled genetic screening in primary immune cells for immunotherapeutic target discovery (Chen et al., 2021; Dong et al., 2019; Gurusamy et al., 2020; Wei et al., 2019; Ye et al., 2019).

With the discovery of numerous immune modulators, understanding how these genes function, whether they are in series or in parallel, will dictate how different combinations of targets will be used therapeutically. Systemically elucidating how these immune regulators interact using CRISPR technologies would require multiple crRNAs targeting an equivalent number of genes. However, Cas9 cannot to process its own CRISPR RNAs (crRNAs). Cas9 systems require individual promoters or the presence of internal endoribonuclease sites to drive the expression of multiple crRNAs for multiplexed gene editing (Nissim et al., 2014; Sakuma et al., 2014; Tsai et al., 2014; Xie et al., 2015). This greatly limits the number of genes that can be studied simultaneously within a given cell due to the limited amount of genetic material that can be efficiently packaged within a pseudoviral vector.

Cas12a (also known as Cpf1) is a class II type V CRISPR endonuclease discovered to have gene editing capabilities in mammalian cells (Fonfara et al., 2016; Zetsche et al., 2015). Further work in engineering Cas12a enhanced its gene editing efficiencies to levels close to Cas9 and related homologs when targeting similar genomic sites (Kleinstiver et al., 2019; Liu et al., 2019). Unlike Cas9 endonucleases, the Cas12a endonuclease family also possesses endoribonuclease activity that allows these enzymes to cleave RNAs upstream of their direct repeat consensus sequences (Fonfara *et al*., 2016; Zetsche *et al*., 2015), which enables this family of enzymes to generate multiple mature crRNAs from a concatenated string of guides (Campa et al., 2019; Zetsche et al., 2017). The *Lachnospiraceae bacterium* (Lb) variant of Cas12a (LbCas12a) has been used to perturb multiple genes simultaneously or sequentially in cancer cell lines (Chow et al., 2018; Chow et al., 2019). The LbCas12a enzyme along with AAV vectors were also utilized in knocking in chimeric antigen receptors (CARs) into the genomic loci of human primary T cells while simultaneously knocking out putative checkpoint inhibitors (Dai et al., 2019). We therefore reasoned that the generation of LbCas12a transgenic mice could facilitate studies pertaining to multiple genetic perturbations and the gene interaction networks in primary immune cells.

Here, we created conditionally and constitutively active LbCas12a knock-in transgenic mice, with multiple lines of founders, to enable simultaneous and efficient multiplexed genetic manipulation in primary immune cells. We demonstrated the utility of these mice on both DNA-level gene editing and protein-level functional knockdown. More importantly, we demonstrated the versatility of genome editing across multiple primary immune cell types including primary CD4 and CD8 T cells, B cells, and bone-marrow derived dendritic cells (BMDCs). These transgenic animals along with the accompanying viral vectors provide a versatile toolkit for *ex vivo* and *in vivo* gene editing for fundamental immunological research, immune engineering, and immunotherapy target discovery.

## Results

### Successful generation of LbCas12a transgenic mice

Cas12a family proteins are large endonucleases with sizes ranging from 1,200 to 1,500 amino acids (Shmakov et al., 2015). Thus, delivery of the complete CRISPR-Cas12a system by viral vectors can be challenging, since packaging large transgenes into viral vectors is problematic and leads to low viral titers. Although delivery of Cas12a proteins in the form of protein/RNP or mRNA is possible, the expression of Cas12a is transient, leading to high variations in editing efficiency. While high variability can be tolerated in experiments targeting a single gene, it could cause unforeseen biases and reproducibility issues with complex CRISPR screens. Therefore, to streamline multiplexed immune cell genome engineering, we developed LbCas12a-transgenic mice with the goal of significantly improving the efficiency and simplicity of multiplexed gene editing in primary immune cells.

We first created a codon optimized LbCas12a knock-in construct by cloning the NLS-LbCas12a-NLS-HA-2A-GFP transgenes into the Ai9 Rosa26-targeting construct to direct transgene recombination between exon 1 and exon 2 of the Rosa26 locus on chromosome 6 (**Fig. 1A**). LbCas12a expression is driven by a CAG promoter but interrupted by a downstream LoxP-3x PolyA Stop-LoxP (LSL) cassette, which allows tissue-specific gene engineering (**Fig. 1A**). LbCas12a gene was flanked by 2 nuclear localization signals (NLSs) to attain better editing efficiency from higher nuclear translocation rates of LbCas12a. Affinity tag 3xHA was fused to the C-terminus of LbCas12a that is immediately followed by a 2A self-cleavage peptide and subsequent fluorescent marker, enhanced GFP (eGFP). The HA tag and eGFP allow detection of LbCas12a by standard molecular and cellular approaches such as western blot and fluorescent microscopy, respectively (**Fig. 1A**). After sequence verification, the LbCas12a Rosa-26-targeting construct was co-administered with SpCas9:Rosa26-crRNA ribonucleoprotein (RNP) into C57BL6/J embryos to generate a LbCas12a knock-in transgenic mouse strain (Rosa26-LSL-LbCas12a). Genotyping and Sanger sequencing verified the successful knock-in of the transgene into the Rosa26 locus.

**Figure 1.**
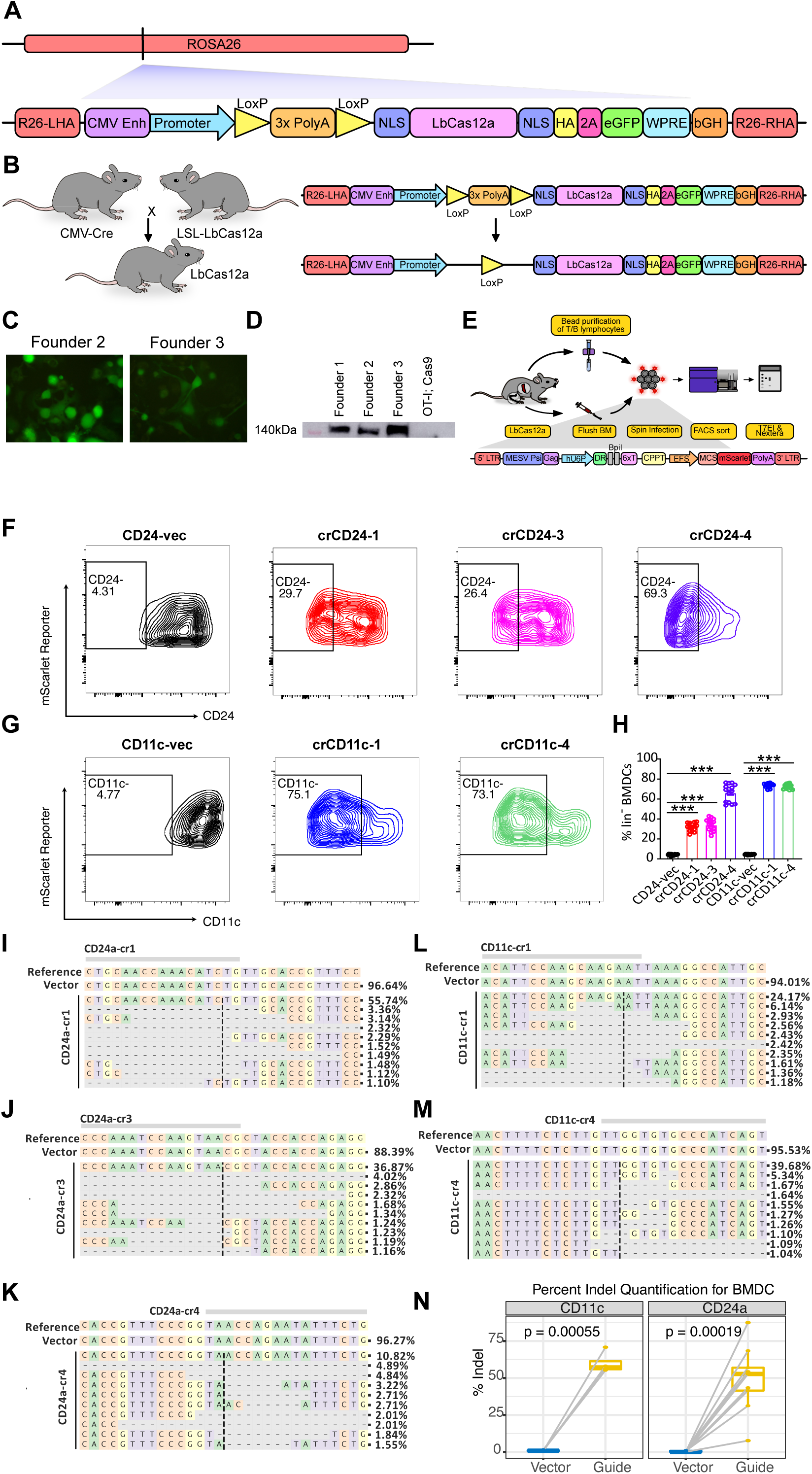
Generation of LbCas12a-transgenic mice and gene editing in BMDCs. (A) Schematic of the LSL-LbCas12a Rosa26-targeting construct. The backbone is Ai9 Rosa26 targeting vector. LbCas12a, labeled with HA tag and enhanced GFP, is expressed under CAG promoter. LoxP-(Stop)3XPolyA-LoxP (LSL) allows Cre-dependent conditional expression of LbCas12a protein. (B) Generation of the constitutively active LbCas12a transgenic mice line (Rosa26-LbCas12a) by crossing the LSL-LbCas12a mouse with CMV-Cre mouse. The LSL-cassette was excised by Cre, leading to constitutively expressing LbCas12a. (C) Fluorescent microscopy showing the expression of LbCas12a-2A-GFP from different founder mice. (D) Western blot showing the expression of LbCas12a protein from different founder mice. The protein is detected by HA tag antibody. OT-I;SpCas9 mice is the negative control. (E) Experimental schematic of the workflow for gene editing with LbCas12a mice in primary immune cell. Primary immune cells are isolated either through beads selection (T/B lymphocytes) or bone marrow flushing (BMDCs). Primary immune cells are then transduced by retrovirus (expression of LbCas12a DR and guide is driven by human U6 promoter). Gene editing efficiency is characterized by flow cytometry, T7E1 assay, and NGS. (F) Flow cytometry analysis on BMDCs’ CD24 with cr1, cr3, and cr4. (G) Flow cytometry analysis on BMDCs’ CD11c with cr1 and cr4. (H) Quantification of CD24 or CD11c negative BMDCs percentage for the guides in E-F, comparing the vector control. (I-K) Allele frequency table from the Nextera sequencing of CD24 guides (L-M) Allele frequency table from the Nextera sequencing of CD11c guides (N) Percent indel quantification of each guide targeting CD24 and CD11c, compared with its corresponding vector control (represented by two dots connected by grey line). In the box plots, mean from guide group and vector control group are compared by paired two-sided t-test and p-values are displayed on the plots. Multiple guides (crRNAs) are pooled and plotted together.

To generate a constitutively active LbCas12a transgenic mice line (Rosa26-LbCas12a), Rosa26-LSL-LbCas12a transgenic mice were then crossed with CMV-Cre mice (**Fig. 1B**). We were able to detect the expression of GFP from primary fibroblasts isolated from Rosa26-LbCas12a mice, and LbCas12a protein in mouse protein lysate by using an anti-HA-tag antibody for western blot analysis (**Fig. 1C-D**). Overall, these data demonstrated the successful generation of Rosa26-LSL-LbCas12a and Rosa26-LbCas12a transgenic mouse lines and expression of LbCas12a protein in the tissue of Rosa26-LbCas12a mice.

### LbCas12a mice enables genetic perturbation in BMDCs and B cells

Current Cas12a crRNA design algorithms are based on either minimizing off-target scores (RGEN) (Park and Bae, 2018), or designed specifically for the *Acidaminococcus sp.* variant of Cas12a (AsCas12a) (CRISPick) (DeWeirdt et al., 2021; Kim et al., 2018), which has been shown to bind to a broader array of crRNA scaffolds compared to LbCas12a (Zhong et al., 2017). To date, there are no suitable algorithms designed for highly functional LbCas12a crRNA design. Nevertheless, we had found that Broad’s CRISPick algorithm designed for AsCas12a can increase the frequency of functional LbCas12a crRNAs (DeWeirdt *et al*., 2021; Kim *et al*., 2018). To identify functional guides (crRNA spacers), we tested the aforementioned algorithms by subcloning the predicted crRNAs targeting lineage-defining immune surface markers into lentiviral plasmids (Sanjana et al., 2014). A murine non-small cell lung cancer cell line expressing LbCas12a transgene (KPD-LbCas12a cell line) was infected by individual lentiviruses, encoding different crRNAs, followed by puromycin selection. Then, genomic DNA was extracted for T7E1 surveyor assay to quantify the cutting efficiency of each guide (**Fig. S1A**). Although originally designed for predicting crRNAs for AsCas12a system, 16 out of 38 total crRNAs (42.1%) predicted by the CRISPick algorithm can mediate target gene editing. On the other hand, only 3 out of 51 crRNAs (5.88%) designed by RGEN led to target genes editing. CRISPick-designed crRNAs were further filtered and enriched by selecting crRNAs with 40% or greater predicted cutting efficiency in their “Target cut %” feature, which we named “CRISPick best cutters”. This resulted in 35 out of 48 crRNAs (72.9%) demonstrating successful gene editing (**Fig. S1B**). As is shown by T7E1 surveyor assay, the average cutting efficiency for crRNAs after filtering is 21.3%, 2.49-fold higher than that of crRNAs before filtering (8.64%) with a p-value 0.00244 (**Fig. S1C**). These data demonstrated that CRISPick algorithm and its “Target cut %” score is a strong indicator for *in vitro* gene editing with LbCas12a. Thus, we used CRISPick for LbCas12a crRNA design and selected crRNAs which demonstrated editing in the KPD-LbCas12a cell line for downstream validation.

To validate that the transgenic LbCas12a mice can be a functional platform for multiplex genome engineering, we tested gene editing in various primary immune cells (BMDCs, B cells, CD4 T cells, and CD8 T cells) isolated from Rosa26-LbCas12a mice. We developed a new retroviral system to further facilitate efficient immune gene editing in conjunction with these transgenic mice. The base vector of this retroviral system (pRetro-hU6-crRNA/BpiI-EFS-mScarlet) utilized a single hU6 promoter to express an individual LbCas12a crRNA or an array of LbCas12a crRNA for multiplexed gene editing upon transduction of mouse primary immune cells. Cloning of crRNAs downstream of the hU6 promoter is mediated by a dual BpiI cloning site, which can be performed with a single cloning step (**Fig. 1E**). Retroviral transduction can be detected by the expression of mScarlet reporter in B cells (CD19^+^), bone marrow-derived dendritic cells (BMDCs) (MHCII^+^CD11b^+^CD11c^+^), and CD4 and CD8 T cells (CD3+) (**Fig. S2A**). Editing efficiency for each guide was assessed by flow cytometry, T7E1 surveyor assay, and Nextera sequencing (**Fig. 1E**). Gene perturbation of highly expressed surface markers in BMDCs, like CD24 and CD11c, can be assessed by measuring the frequency of reporter-positive cells that downregulated the expression of targeted protein. Flow cytometry data showed that crCD24-1, crCD24-3, and crCD24-4 resulted in 29.7%, 26.4%, and 69.3% knockdown of CD24 respectively, while crCD11c-1 and crCD11c-4 yielded 63.1% and 61.8% knockdown of CD11c in BMDCs (**Fig. 1F, Fig. 1G, Fig. 1H**). Quantification gate was set up such that less than 5% cells knockdown in the vector control group (**Fig. 1F, Fig. 1G, Methods**). crCD24-4 was also the most efficient guide among the three by T7E1 surveyor assay quantification, similar to crCD11c-1 (**Fig. S1D**). Nextera libraries were generated for each sample and submitted for Next Generation Sequencing (NGS). Nextera sequencing results corresponded well with the flow cytometry and T7E1 surveyor data, with crCD24-4 being the best crRNA targeting CD24 (resulting up to 87.5 % indels) and crCD11c-1 being the slightly better crRNA for CD11c (generating up to 70.8% indels) (**Fig. 1I-N**). Allele frequency plots demonstrated that deletion was more prominent than insertion and the mutations were roughly centered at the predicted cutting site of LbCas12a (**Fig. 1I-M**). These data showed successful highly efficient gene editing and protein knockdown of specific surface markers in BMDCs derived from Rosa26-LbCas12a mice.

We also tested whether these transgenic mice and the retroviral system can be utilized for B cell gene editing. B cells were isolated from LbCas12a mice and transduced by retrovirus. DNA-level gene editing as well as protein-level knockdown were analyzed (**Fig. 1E**). The relative fold difference in B220 expression between vector-treated B cells and crB220-treated B cells (surface protein knockdown), measured by flow cytometry, spanned from 6.25% (crB220-1) to 67.6% (crB220-3) (**Fig. S3A**). Gene editing of B220 by the three guides was also measured by T7E1 assay (**Fig. S3B**). Nextera sequencing result showed that crB220-3 is the most efficient crRNA (16.1% editing efficiency), while crB220-1 and crB220-2 led to 2.3% and 9.6% indel (**Fig. S3C-E, Fig. S3H**). For CD30, two crRNAs, crCD30-1 and crCD30-2 resulted in 13.1% and 2.8% indel in B cell (**Fig. S3F-H**). These data show successful gene editing and protein knockdown of specific surface markers, although at relatively lower efficiency, in B cells derived from Rosa26-LbCas12a mice.

### LbCas12a mice enables genetic perturbation in CD4 and CD8 T cells

We then tested gene editing in CD4 and CD8 T cells derived from LbCas12a mice using the same system. Some of the crRNAs were designed to target surface proteins which have similar expression level in CD4 and CD8 T cells (Thy1, Pd-1, Ctla-4), while others were targeting lineage specific surface proteins (CD4, CD8a). In CD4 T cells, Thy1 and Cxcr4 were knocked down by 40.6% and 68.7% with the top crRNA, as measured by flow cytometry (**Fig. 2A, Fig. 2B, Fig. 2D**). Results in crCD4-1 group showed that 74.2% of the CD4 T cells lost their CD4 expression, compared to only 16.7% cells in the vector control group were negative. (**Fig. 2C, Fig. 2D**). As for CD8 T cells, crCD8a-2 is the best crRNA targeting CD8a, achieving 71.4% knockdown efficiency as indicated by the flow cytometry analysis (**Fig. 2E, Fig. 2G**). crRNAs targeting Thy1 also achieved knockdown efficiency around 30% by flow cytometry analysis (**Fig. 2F, Fig. 2G**).

**Figure 2.**
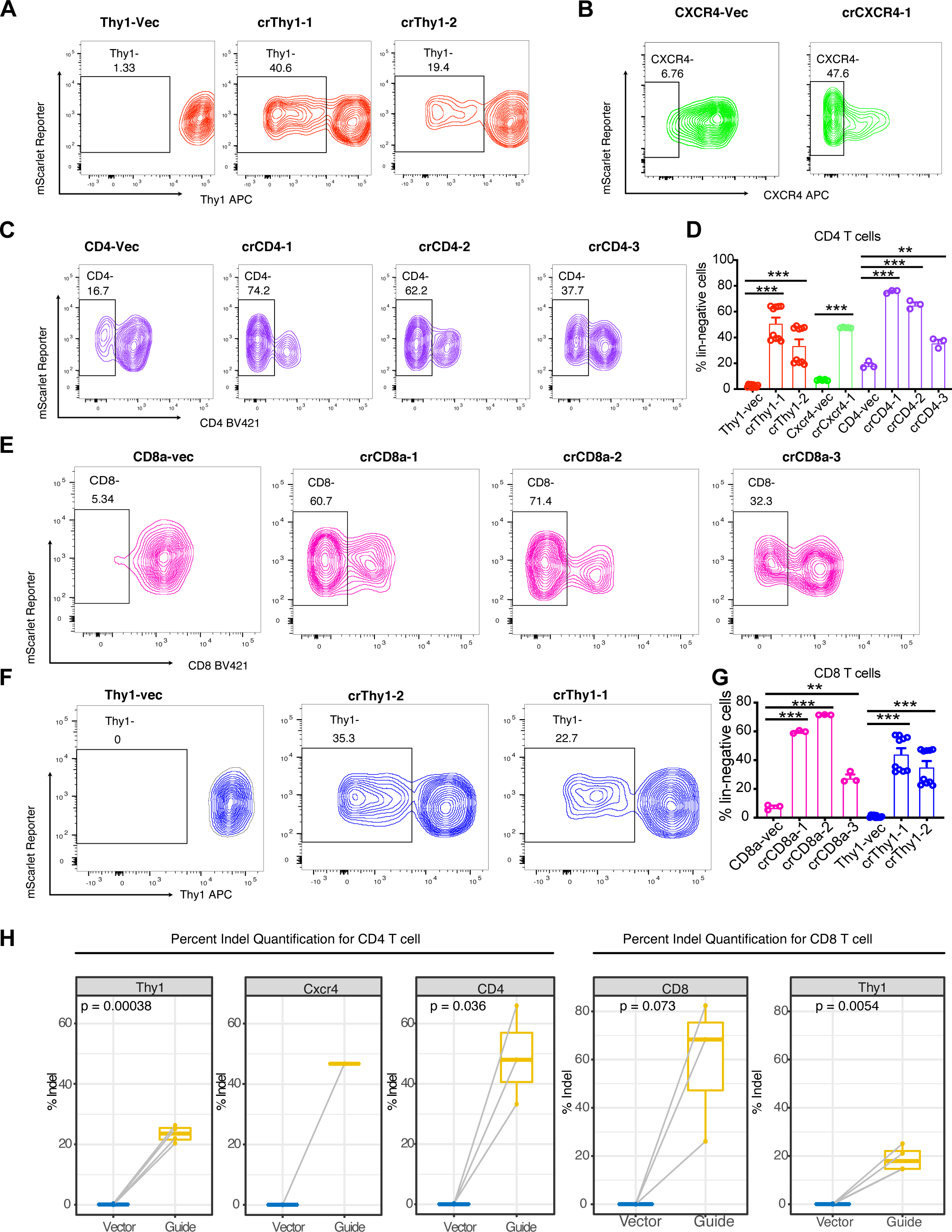
Analysis of gene perturbation in CD4 and CD8 T cells with LbCas12a mice. (A) Flow cytometry analysis on CD4 T cells’ Thy1 with cr1 and cr2. (B) Flow cytometry analysis on CD4 T cells’ Cxcr4 with cr1. (C) Flow cytometry analysis on CD4 T cells’ CD4 with cr1, cr2, and cr3. (D) Quantification showing the percentage of CD4 T cells that downregulate Thy1, Cxcr4, and CD4 protein expression when transduced with the indicated crRNA. For bar plots, data are shown as mean ± s.e.m. * = p < 0.05, ** = p < 0.01, *** = p < 0.001, by unpaired two-sided t-test. (E) Flow cytometry analysis on CD8 T cells’ CD8a with cr1, cr2, and cr3. (F) Flow cytometry analysis on CD8 T cells’ Thy1 with cr1 and cr2. (G) Quantification showing the percentage of CD8 T cells that downregulate CD8a and Thy1 protein expression when transduced with the indicated crRNA. For bar plots, data are shown as mean ± s.e.m. * = p < 0.05, ** = p < 0.01, *** = p < 0.001, by unpaired two-sided t-test. (H) Percent Indel quantification of each guide targeting *Thy1*, *Cxcr4*, CD4/8 in CD4 and CD8 T cells, compared with its corresponding vector control (represented by a paired of dots connected by a grey line). In the box plots, mean from guide group and vector control group are compared by paired two-sided t-test and p-values are displayed on the plots. Multiple guides (crRNAs) are pooled and plotted together.

Nextera library preparation followed by Next Generation Sequencing was used to quantify the gene editing efficiency on molecular level. For all crRNAs targeting surface proteins in CD4 and CD8 T cells, higher indel percentages were observed compared to vector controls (**Fig. 2H**). crCD4-1 generated the highest (66%) indels for the CD4 T cells (**Fig. 2H, Fig. S4A-M**). Similarly, crCD8a-1 generated the highest (82%) indels for CD8 T cells (**Fig. 2H, Fig. S5A-M**). We suspected the differential knockout efficiency between different target genes was due to difference in their chromatin accessibility, as highly expressed genes had higher gene editing efficiency. We then went on to test this hypothesis by targeting Cxcr4 and Il7ra, which were found to be expressed higher in cultured CD4 T cells compared to CD8 T cells (**Fig. S2B-C, Fig. S2E-F**). Higher editing efficiency, as shown by flow cytometry (**Fig. SC, Fig. S2F**) and T7EI assay (**Fig. S2D, Fig. S2G**), was observed in CD4 T cells.

### Double targeting of immune cells surface proteins using single crRNA arrays in primary cells from LbCas12a mice

An important objective of the LbCas12a transgenic mice is to facilitate multiplex genome editing on single cell level. Because T7E1 surveyor assay and Next Generation Sequencing of bulk cells survey the genetic editing from a mixed population of cells, flow cytometry was used to quantify the ability of our system to perturb multiple genes simultaneously on single cell level. Two pairs of highly efficient, functional crRNAs (crCD24-3; crCD43-1 and crCD24-3; crCD11c-1) were selected and concatenated into crRNA arrays. LbCas12a BMDCs were then transduced with retroviruses harboring either vector control, single crRNAs (crCD24-3, crCD43-1, and crCD11c-1), or double crRNAs (crCD24-3; crCD43-1 and crCD11c-1; crCD24-3).

For the first pair of crRNAs (crCD24; crCD43), flow cytometry analysis indicated that single crCD24 and crCD43 result in 54.08% and 12.1% of cell with knockout, respectively. Double crRNA array crCD24;crCD43 knocked out CD24 and CD43 at a similar level to the single crRNA counterparts (52% for CD24 and 14.51% for CD43) (**Fig. 3A-3B**). Interestingly, 11.8% of the cells had both CD24 and CD43 knockout, which was observed uniquely in the double crRNA group (**Fig. 3A-B**). The second crRNA array (crCD24; crCD11c) demonstrated higher percentage (26.4%) of single cell level double knockout (**Fig. 3C-D**), whereas crCD24 and crCD11c single crRNAs only knocked down specific corresponding single protein but not the other untargeted protein (**Fig. 3C-D**). To confirm the lower expression level was indeed caused by genome editing, Next Generation Sequencing was used to survey the indel frequency at the molecular level. For the crRNA array (crCD24; crCD43), over 70% and 65% of the indels were on *CD24a* and *CD43* gene, respectively (**Fig. 3E-F**). Indel frequency was closed to zero in the vector control (**Fig. 3E-F)**. Collectively, our data showed the multiplex capability and versatility of our transgenic mice and retroviral system for genome engineering. These transgenic animals along with our retroviral vectors provide a versatile toolkit for a broad range of applications of *ex vivo* and *in vivo* gene editing, especially for immunology and gene discovery.

**Figure 3.**
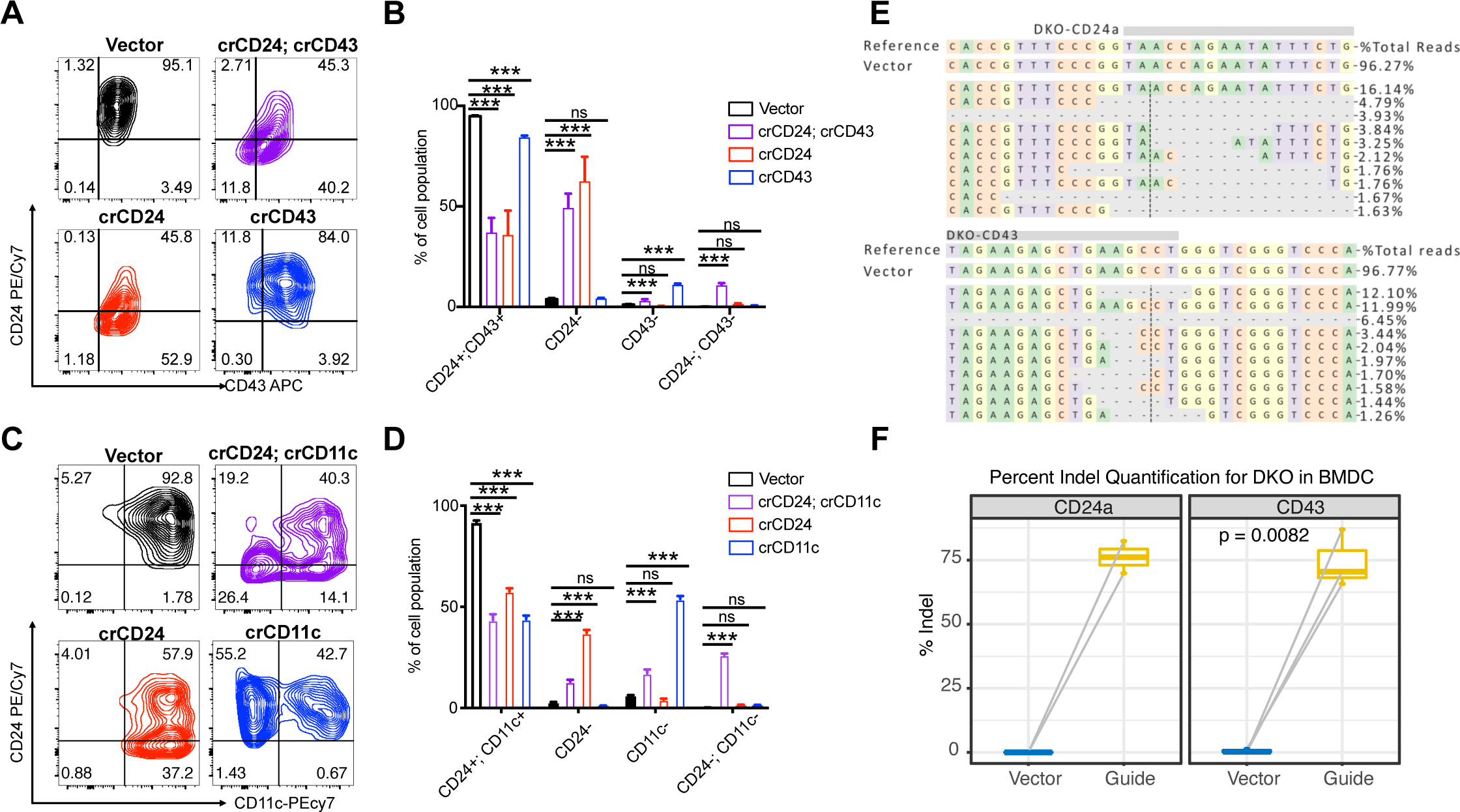
Double knockout in BMDCs from LbCas12a Transgenic Mice. (A) Representative contour plots of Reporter+ BMDCs showing CD24 expression in relation of CD43 expression for indicated samples. CD43-APC and CD24-PE/Cy7 are used for the flow staining. crRNA array that contains both crCD24 and crCD43 is compared with single crRNA or vector control. (B) Quantification showing the percentage of BMDCs that downregulate CD24, CD43, or both for each indicated sample. For bar plots, data are shown as mean ± s.e.m. * = p < 0.05, ** = p < 0.01, *** = p < 0.001, by unpaired two-sided t-test. (C) Representative contour plots of Reporter+ BMDCs showing CD24 expression in relation of CD11c expression for indicated samples. CD24-APC and CD11c-PE/Cy7 are used for the flow staining. Double crRNA array that contains both crCD24 and crCD11c is compared with the single crRNA or vector control. (D) Quantification showing the percentage of BMDCs that downregulate CD24, CD11c, or both for each indicated sample. For bar plots, data are shown as mean ± s.e.m. * = p < 0.05, ** = p < 0.01, *** = p < 0.001, by unpaired two-sided t-test. (E) Allele frequency table from the Nextera sequencing of double crRNA array. (F) Percent Indel quantification of the double crRNA array targeting both CD24a and CD43 in BMDCs, compared with its corresponding vector control (represented by two dots connected by grey line). In the box plots, mean from guide group and vector control group are compared by paired two-sided t-test and p-values are displayed on the plots. Multiple guides (crRNAs) are pooled and plotted together.

## Discussion

Discovery of genetic regulators is critical for basic immunology and the development of immunotherapy for cancer and autoimmune disorders. CRISPR-based gene editing and screening have accelerated the pace of such discovery, especially with the innovation of CRISPR animal models. Various transgenic animals have been generated, primarily centered on the SpCas9 effector and its variants. However, due to its inability to process its own CRISPR RNA (crRNA), simultaneous editing of multiple genes using the Cas9 system is challenging especially due to the packaging constraints of viral vectors. Multiplexed gene editing using Cas9 systems would require a unique U6 promoter for each crRNA, a crRNA with an average length of 100bps, and a multistep cloning process to express each crRNA. In contrast, Cas12a/Cpf1 is capable of processing its own crRNAs using its endoribonuclease activity from a concatenated string of crRNAs that is driven by a single promoter. More importantly, each crRNA is encoded by only 40-44 bps, which is 2.5-times smaller than a Cas9 crRNA. Therefore, simplification of the delivery of the Cas12a/Cpf1 machinery can streamline genome editing, genetic discovery applications, and complex gene network analyses.

LbCas12a knock-in mice and accompanied retroviral vector system can serve as a versatile new set of genetic tools. The simplicity of multiplexing crRNAs into a single crRNA array streamlines the cloning process and investigations of multiplexed gene perturbations. Using these technologies, we demonstrated that these mice enable *bona fide* efficient genome editing and protein knockdown across multiple types of immune cells, reaching efficiencies as high as 70-80%. In addition, we have demonstrated highly efficient, multiplexed gene-editing within a single primary immune cell. While we focus on *ex vivo* applications of primary immune cells, various applications of *in vivo* multiplexed gene editing are in principle also feasible using these mice. Beyond our demonstration of immune engineering, the application of these mice can be extended to many other fields such as cell biology, neuroscience, developmental biology, cancer biology, and drug target discovery. Importantly, with the simplicity of multiplexed gene targeting with single crRNA arrays, LbCas12a mice can serve as an important tool at elucidating how genes interact in complex networks.

The generation of Cas9-mice and its many variations has greatly served the scientific community, enabling gene editing and CRISPR screens in primary immune cells (Chen *et al*., 2021; Dong *et al*., 2019; Gurusamy *et al*., 2020; Platt *et al*., 2014; Wei *et al*., 2019; Ye *et al*., 2019). Nevertheless, these studies are limited to mostly single gene perturbations because of the limitation of Cas9. The LbCas12a knock-in mice are orthogonal resources, which not only expand the complexity of multiplex gene perturbations, but also enable new multi-dimensional genetic perturbation studies as it can easily be crossed to other Cas9- or dCas9 mice (Zhou *et al*., 2018), as well as various other transgenic strains. The Cas12a mice enables future applications that utilizes compound transgenic lines, such as Cas12a;Cas9 or Cas12a;dCas9 mice, to enable simultaneous and orthogonal gene activation and gene perturbation studies due to the unique structure of the LbCas12a and SpCas9 crRNAs. Collectively, the recently created LbCas12a knock-in mice and associated tools here provide a versatile platform that enables efficient multiplexed immune cell engineering, as well as a broad range of potential applications in basic biomedicine and therapeutic target discovery.

## Acknowledgments

We thank all members of the Chen laboratory, as well as our colleagues in Department of Genetics, Systems Biology Institute, Cancer Systems Biology Center, MCGD Program, Immunobiology Program, BBS Program, Yale Cancer Center, Yale Stem Cell Center, RNA Center and Center for Biomedical Data Sciences at Yale for assistance and/or discussion. We thank the Yale Genome Editing Center, Yale Center for Genome Analysis, Yale Center for Molecular Discovery, High Performance Computing Center, West Campus Analytical Chemistry Core, and Keck Biotechnology Resource Laboratory at Yale, for technical support.

S.C. is supported by NIH/NCI/NIDA (DP2CA238295, R01CA231112, U54CA209992-8697, R33CA225498, RF1DA048811), DoD (W81XWH-17-1-0235, W81XWH-20-1-0072, W81XWH-21-1-0514), Damon Runyon Dale Frey Award (DFS-13-15), Melanoma Research Alliance (412806, 16-003524), St-Baldrick’s Foundation (426685), Breast Cancer Alliance, Cancer Research Institute (CLIP), AACR (17-20-01-CHEN), The Mary Kay Foundation (017-81), The V Foundation (V2017-022), Alliance for Cancer Gene Therapy, Sontag Foundation (DSA), Pershing Square Sohn Cancer Research Alliance, Dexter Lu, Ludwig Family Foundation, Blavatnik Family Foundation, and Chenevert Family Foundation. MBD is supported by NIH MSTP training grant (T32GM007205). RC is supported by NIH MSTP training grant (T32GM007205) and NRSA fellowship (F30CA250249). XD is supported by Charles H. Revson Senior Postdoctoral Fellowship.

## STAR Methods

### Institutional Approval

This study has received institutional regulatory approval. All recombinant DNA and biosafety work were performed under the guidelines of Yale Environment, Health and Safety (EHS) Committee with an approved protocol (Chen-rDNA 15-45; 18-45; 21-45). All animal work was performed under the guidelines of Yale University Institutional Animal Care and Use Committee (IACUC) with approved protocols (Chen 2018-20068; 2021-20068).

### crRNA Design and Cloning

The design of LbCas12a crRNAs primarily utilizes Broad’s CRISPick algorithm (DeWeirdt *et al*., 2021; Kim *et al*., 2018). To further enrich for highly functional LbCas12a crRNAs, CRISPick-designed guides were chosen based on their predicted ability to cut with greater than 40% efficiency. Guides were then subcloned into lentiviral and retroviral plasmids, as previously described (Sanjana *et al*., 2014). Designs of lentiviral and retroviral plasmids are described below.

### Lentiviral Production

pLenti-U6-DR-crRNA(Esp3I)-EF1a-Puro-P2A-Firefly luciferase transfer plasmid was generated as previously described (Chow *et al*., 2019). To generate lentivirus, 10^6^ HEK293 cells (Invitrogen) were plated into 6-well plate the night before in D10 media (DMEM + 10% FBS + 1% Pen/Strep). The next day, media were changed, and cells were allowed to incubate at 1-2hr at 37°C prior to transfection. 1μg of transfer plasmid, 0.75μg of psPAX2 packaging plasmid, and 0.5μg of pMD2.G envelop plasmid were added to 50μl of DMEM media. In a separate tube, 3μl of LipoD293 (Signagen) was added to 50μl of DMEM media and then quickly added to DNA:DMEM mixture. Transfection mixture was then incubated at room temperature (RT) for 10-15 minutes before being added to the HEK293 cells. Lentivirus in the supernatant was collected 48 hours after transfection. Spun down at 3000xg for 5 minutes and then filtered through 0.22μm filter. Lentivirus was either used directly for experimentation or stored in −80°C before use.

### Cell Lines and Lentiviral Infection

Non-small cell lung cancer cell line expressing LbCas12a (KPD-LbCas12a) was generated and cultured as previously described (Chow *et al*., 2019). The day before lentiviral transduction, 3 x 10^5^ KPD-LbCas12a cells were plated in 6-well plate. Uninfected KPD-LbCas12a cells were included on each plate to determine extent of Puromycin positive selection. Vector controls were included for each experiment to calculate the cutting efficiency of each crRNA. Media were changed and supplemented with 8μg/ml polybrene (Millipore) to increase transduction efficiency. 1ml of lentivirus was added to indicated well and incubated overnight at 37°C. Media was changed the following morning and supplemented with 5μg/ml Puromycin (Gibco). Puromycin selection was continued until uninfected KPD-LbCas12a cells were no longer viable as determined by light microscopy. Upon the completion of puromycin selection, infected KPD-LbCas12a cells were trypsonized and collected. Cell pellets were washed twice with PBS prior to genomic DNA extraction and purification. QIAamp DNA Mini Kit was used to extract and purify genomic DNA of infected cells.

### Determine Gene Editing Efficiency

To determine the cutting efficiency of each crRNA, genomic sequence of each gene was obtained from Ensembl genome browser. Sequences 1.5kb both upstream and downstream of the crRNA target sequence were input into NCBI Primer Blast tool. *Mus musculus* genome was used to identify unique primers within the mouse genome. Surveyor PCR primers were designed such that the difference between the right side of crRNA target sequence and the left side would be greater than 100nt and no fragment length should be less than 100nt, the lower limit the low range quantitative DNA ladder (Invitrogen). PCR products were restricted to 600-1300nt with preferences for smaller PCR fragmentation. Purified PCR product was subjected to T7 endonuclease 1 digestion (NEB) and Nextera sequencing according to Illumina protocol to quantify gene-editing efficiency.

### Mice

Cre-dependent LbCas12a mice were generated by pronuclear injection of transgenic expression cassette, IDT Alt-R® HiFi Cas9 Nuclease V3, and Rosa26-targeting crRNA (5’-ACTCCAGTCTTTCTAGAAGA-3’) (Chu et al., 2015). Codon optimized LbCas12a cDNA that subcloned into the Ai9 Rosa26-targeting vector, giving rise to LSL-NLS-LbCas12a-NLS-3x HA-P2A-GFP transgenic expression cassette (Madisen et al., 2010; Sanjana *et al*., 2014). Unique sequencing and genotyping primers were designed using NCBI Primer Blast tool against *Mus musculus* genome. Transgenic mice were first identified using internal LbCas12a primers (forward: 5’-TGCTGAGCGATCGGGAGTCT-3’ and reverse: 5’-TGGTCCACCTTCAGCAGGATG-3’) to identify presence of LbCas12a transgene. *Rosa26* primers external to Ai Rosa26-targeting vector were used to amplify and sequence confirm correctly targeted transgenic expression cassette by Sanger sequencing. Constitutive LbCas12a mice were generated by crossing Cre-dependent LbCas12a mice to CMV-Cre mice (Jackson Laboratories). Other mouse strains used for experiments include C57BL6/J (Jackson). Mice, both sexes, between the ages of 6-12 weeks of age were used for the study. All animals were housed in standard individually ventilated, pathogen-free conditions, with 12h:12h or 13h:11h light cycle, room temperature (21-23°C) and 40-60% relative humidity.

### Mouse Fibroblast Cultures

Mouse fibroblast cultures were obtained by digesting mouse ear using Collagenase/Dispase (Roche) for 30 minutes at 37°C. Supernatant was collected and washed with 2% FBS. Cell suspensions were the filtered through 40μm filter and resuspended in D10 media.

### Widefield Fluorescence Microscopy

Images of mouse fibroblast cultures were taken when cells reached 60-75% confluency using Leica DMi8 Widefield Fluorescence Microscope with a Lumencor Spectra X light engine and a 510/25-25 filter.

### Western Blot Analysis

Mouse fibroblasts were trypsonized with TrypLE (Thermo) for 5 minutes at 37°C. Cells were washed once with D10 media followed by 2 serial washes with PBS. Pierce IP Lysis Buffer was added to cell pellets and incubated for 5 minutes on ice. NuPAGE LDS sample buffer with beta-mercaptoethanol was added to cell lysates and heated at 70°C for 10 minutes. Samples were then heated at 95°C for 3 minutes and placed on ice for 1 minute before immunoblotting. 4∼20% Tris-Glycine gels were used for gel electrophoresis. Proteins were then transferred to 0.45μm PVDF membrane. Membranes were blocked with 5% BSA, stained with mouse anti-HA tag antibody (Thermo) overnight at 4°C and stained with goat anti-mouse IgG (H+L) Secondary Antibody, HRP (Thermo) for 1 hour at room temperature. Proteins were detected using Pierce ECL substrate.

### Generation of retroviral-Cas12a-guide vector system

The retroviral base vector, pRetro-U6-DR-crRNA(BpiI)-EFS-mScarlet was generated by subcloning U6-DR-crRNA(Esp3I) regions from pLenti-U6-DR-crRNA(Esp3I)-EF1a-Puro-P2A-Firefly luciferase transfer plasmid and mScarlet from pEB2-mScarlet transfer plasmid (Balleza et al., 2018) into MSCV-IRES-Thy1.1 retroviral transfer plasmid (Wu et al., 2006). Esp3I cut site in pRetro-U6-DR-crRNA(Esp3I)-EFS-mScarlet was mutated to BpiI cut site due to the presence of multiple Esp3I cut sites in resulting plasmid. Various crRNA arrays were cloned into the retroviral base vector.

### Retroviral Production

To generate retrovirus, 4 x 10^7^ HEK293 cells were cultured into 150mm plate the night before in D10 media. The next day, media were changed, and cells were allowed to incubate for 1-2hr at 37°C prior to transfection. 16μg of transfer plasmid and 8μg of pCL-Eco packaging plasmid was added to 1ml of DMEM media. In a separate tube, 62.5μl of LipoD293 was added to 1ml of DMEM media and vortexed prior to being added to the DNA:DMEM mixture. Transfection mixture was then incubated at room temperature (RT) for 10-15 minutes before adding to HEK293 cells. Retrovirus in the supernatant was collected 48 hours after transfection, spun down at 3000g for 5 minutes, and then filtered through 0.22μm filter. Retrovirus was then concentrated by diluting autoclaved 40% PEG8000 (m/v) to final concentration of 8% PEG8000. Viral supernatant was incubated overnight at 4°C. The next morning retrovirus was spun down at 1500g for 30 minutes, supernatant aspirated, and pegylated viral pellet was resuspended with 1ml of fresh complete RPMI (RPMI + 10% FBS + 1% L-glutamine +1% Pen/Strep + 50uM beta-mercaptoethanol). Retrovirus was then stored in −80°C before use.

### Lymphocyte Isolation and Culture

Spleens were isolated from LbCas12a mice and placed in ice-cold 2% FBS [PBS (Gibco) + FBS (Sigma)]. Single cell suspensions were prepared by mashing spleens through a 100um filter. Splenocytes were suspended in 2% FBS. RBCs were lysed with ACK Lysis Buffer (Lonza), incubated for 2 mins at room temperature, and washed with 2% FBS. Lymphocytes were filtered through a 40μm filter and resuspended with MACS Buffer (PBS + 0.5% BSA + 2mM EDTA). CD4+ and CD8+ T cells were isolated using mouse CD4 and CD8 T cell isolation kit from Miltenyi, respectively. B cells were isolated using mouse B cell isolation kit from Miltenyi. All lymphocytes were resuspended to a final concentration of 2 x 10^6^ cells/ml. T cell cultures were cultured in 96-well round bottom plates with plate-bound anti-CD3 (1μg/ml; Biolegend) and soluble anti-CD28 (1μg/ml; Biolegend), IL-2 (2ng/ml; Peprotech), and IL-12 (2.5ng/ml; Peprotech). B cells were cultured in 96-well round bottom plates with plate-bound anti-IgM (1μg/ml; Southern Biotech) and soluble anti-CD40 (1μg/ml; Biolegend), IL-4 (5ng/ml; Peprotech), IL-21 (10ng/ml; Peprotech), and recombinant human BAFF (10ng/ml; R&D).

### BMDC Isolation and Culture

The femur and tibia were isolated from LbCas12a mice and placed in ice-cold 2% FBS [PBS (Gibco) + FBS (Sigma)]. Bones were sterilized by submerging bones in 70% ethanolfor 2 minutes followed by washing the bones 2 times. The ends of the bones were carefully removed with sterilized instruments and the bone marrow was flushed out using 25G needle. RBCs were lysed with ACK Lysis Buffer (Lonza), incubated for 2 mins at room temperature, and washed with complete RPMI. Cells were filtered through a 40μm filter and resuspended with complete RPMI to final concentration of 2 x 10^6^ cells/ml. Media was supplemented with 25ng/ml GM-CSF (Peprotech) and plated in 24-well plate.

### Spin Infection

Final concentration of 8μg/ml polybrene was added cells 18-24 hours after plating. Cells were incubated for 30 minutes at 37°C. Retrovirus was diluted 1:2 with complete RPMI and supplemented to final concentration of 8μg/ml polybrene and 10ng/ml IL-2 for T cells, 10ng of IL-4, IL-21, and hBAFF for B cells, and 25ng/ml GM-CSF for BMDCs. Media were removed from cells and retroviral cocktails were added to indicated cells. Cells were pipetted up and down. Cells were spun down at 2000rpm (∼900g) for 90 minutes at 37°C. Upon completion, cells were incubated for an additional 30 minutes at 37°C before media was completely changed. B cells were continued to be cultured for an additional 5-6 days. T cells were cultured for an additional 7 days. BMDCs were cultured for an additional 8 days. Media was changed every day and cells were split every other day before flow cytometric analysis and sorting.

### Antibody and Flow Cytometry

Generally, cells were then stained with indicated antibody cocktails suspended in MACS Buffer for 30 mins on ice. For chemokine receptors and cycling proteins, cells were stained at 37°C for 30 minutes followed by surface staining for 30 minutes on ice. For BMDCs and B cells, cells were stained with anti-CD16/32 to neutralize IgG Fc receptors prior to any surface protein staining. LIVE/DEAD Fixable Near-IR Dead Cell Stain (Invitrogen) was included in surface stains to exclude dead cells from analysis. B cell lineage markers include anti-CD19 BV421; occasionally anti-CD24 BV421 or PE/Cy7 was used as a B lineage defining surrogate. T cell lineage markers include anti-CD3 PE/Cy7 and anti-CD4/8 BV421; occasionally anti-CD90.2 (Thy1.2) APC was used a T cell lineage surrogate. BMDC lineage markers include anti-MHCII FITC, anti-CD11b BV421, and CD11c PE/Cy7. Infected leukocytes were defined as Lin+mScarlet+. Other antibody stains include anti-CD14 APC, anti-CD24 APC, anti-CD30 APC, anti-CD43 APC, anti-CD96 APC, anti-CD115 APC, anti-CD125 (Il5ra) APC, anti-CD127 (Il7ra) APC, anti-CD152 (Ctla4) APC, anti-CD172A (Sirpa) APC, anti-CD184 (Cxcr4) APC, and anti-CD279 (Pdcd1) APC. Samples were collected sorted on a BD FACSAria cell sorter with 4 lasers (405nm, 488nm, 561nm and 640nm) to > 95% purity. Data was analyzed using FlowJo software 10.8.0 (BD) on a MAC® workstation.

### Genomic DNA extraction

20μl of Epicentre QuickExtract was added for every 50,000 cells sorted. DNA from sorted leukocytes was extracted by incubating cells for 65°C for 30 minutes, and subsequently incubated at 95°C for 5 minutes. Before setting up surveyor PCR reaction, samples were rigorous vortexed to homogenize DNA.

### Nextera library preparation and sequencing

PCR products from the T7E1 essay for each sample were tagged, amplified, and barcoded using Nextera XT DNA Library Prep Kit (Illumina). The library for each sample was quality controlled and quantified separately using the 4150 TapeStation System (Agilent), followed by library pooling and PCR clean up using QIAquick PCR Purification Kit (Qiagen). Another round of quality control and quantification was conducted by running on the 4150 TapeStation System (Agilent). Libraries were denatured and diluted to 10 pM according to Denature and Dilute Libraries Guide (Illumina), before loading on MiSeq (Illumina) for sequencing.

### Next Generation Sequencing data processing and quantify gene modification percentage

FASTQ reads were quality controlled by running FastQC v0.11.9 (Andrews, 2010) and contaminations by Nextera transposase sequence at 3’ end of reads were trimmed using Cutadapt v3.2 (Martin, 2011). Processed reads were aligned to amplicon sequence and quantified for insertions, deletions (indel), and substitution using CRISPResso2 v2.1.3 (Clement et al., 2019). Specifically, we retrieved amplicon sequences, which were 150 ∼ 250 bp (according to the length of reads) flanking crRNA target sites, from the mm10 genome. A 5-bp window, centered by predicted LbCas12a cutting sites, was used to quantify genetic modification for each crRNA in both vector control groups and experimental groups (-w 5 -wc -2). Allele frequency plots were generated with CRISPResso2 (--annotate_wildtype_allele WT --plot_window_size 12). Percent-modification data from each sample were aggregated for analysis and visualization in R. For quality control purpose, guide-vector pairs for sample 20210625-BMDC-CD24a-cr3 and 20210623-BMDC-CD43-2×5 were filtered out due to high background mutation frequency in vector control.

### Sample size determination

Sample size was determined according to the lab’s prior work or from published studies of similar scope within the appropriate fields.

### Replication

Number of biological replicates (usually n >= 3) are indicated in the figure legends. Key findings (non-NGS) were replicated in at least two independent experiments. NGS experiments were performed with biological replications as indicated in the manuscript.

### Randomization and blinding statements

Regular *in vitro* experiments were not randomized or blinded. High-throughput experiments and analyses were blinded by barcoded metadata.

### Standard statistical analysis

Standard statistical analyses were performed using regular statistical methods. GraphPad Prism, Excel and R were used for all analyses. Different levels of statistical significance were accessed based on specific p values and type I error cutoffs (e.g., 0.05, 0.001, 0.0001). Further details of statistical tests were provided in figure legends and/or supplemental information.

### Code availability

The code used for data analysis and the generation of figures related to this study are available from the corresponding author upon reasonable request.

### Data and resource availability

All data and analyses for this this study are included in this article and its supplementary information files. Processed data for NGS or omics data are provided in Supplemental Datasets. Raw sequencing data will be deposited to NIH Sequence Read Archive (SRA) or Gene Expression Omnibus (GEO). Various materials are available at commercial sources listed in Key Resources Table (KRT). Transgenic mice, vectors, cell lines, other relevant information, or data unique to this study are available from the corresponding author upon reasonable request.

## Supplemental Figures

**Figure S1.**
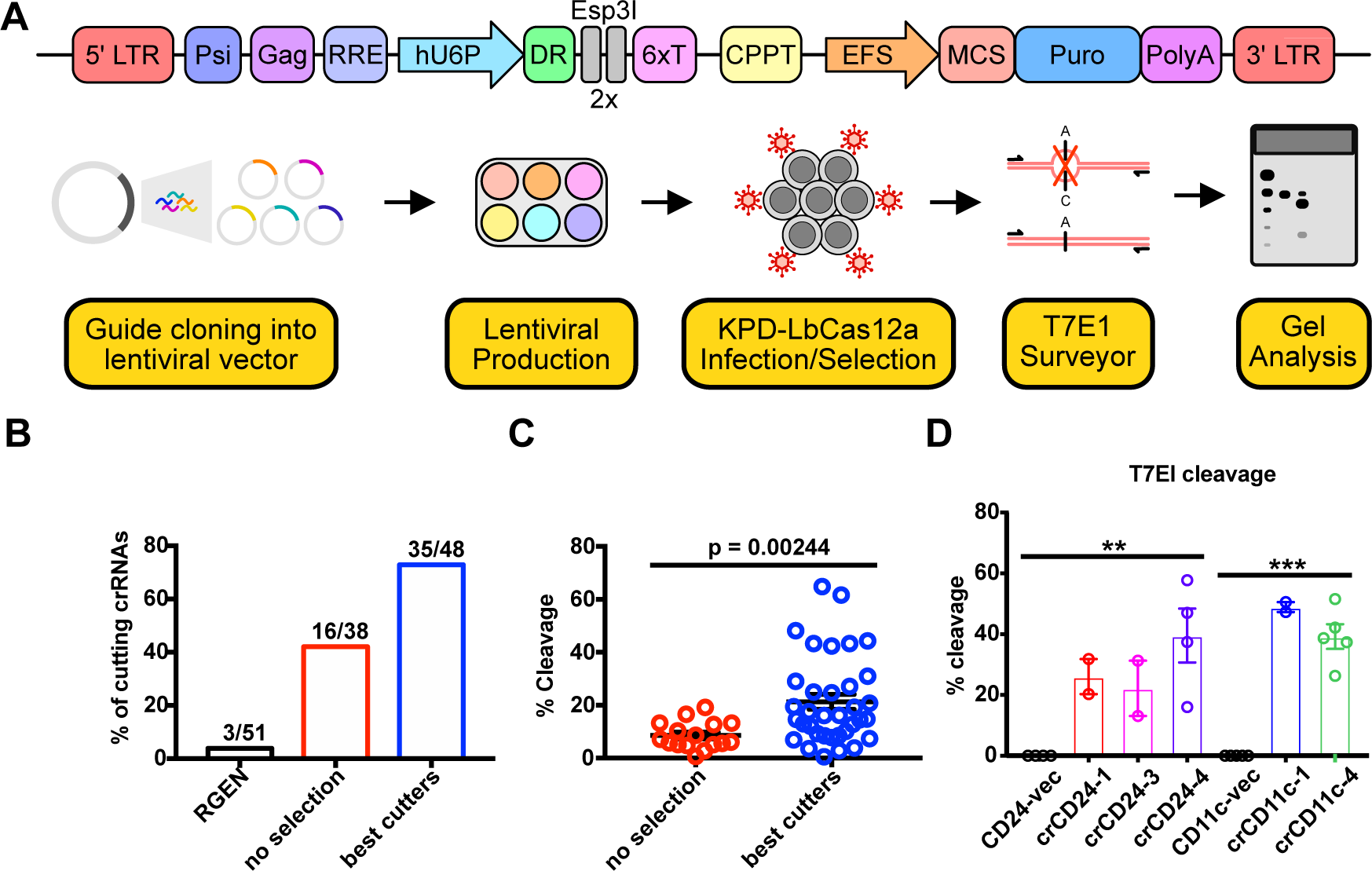
Guide testing in small-cell lung cancer cell line and BMDCs. (A) Schematic of the experimental workflow for testing guides in KPD-LbCas12a cell line. A lentiviral vector, which express crRNA with hU6 promoter is used. Puromycin selection for the infected cells is conducted before the T7E1 assay to characterize the cutting efficiency. (B) Percentage of functional LbCas12a guides based on the crRNA-designing algorithms: RGEN, CRISPick with no selection, and CRISPick with selection. (C) Dot plot showing the cutting efficiency (in percent cleavage) of guides tested in KPD-LbCas12a cell line using unmodified and modified CRISPick algorithm. Cutting efficiency was determine by T7E1 endonuclease surveyor assay. (D) Quantification of T7E1 assay for guides that knock out CD24a and CD11c in BMDCs. For dot-bar plots, data are shown as mean ± s.e.m. * = p < 0.05, ** = p < 0.01, *** = p < 0.001, by unpaired two-sided t-test.

**Figure S2.**
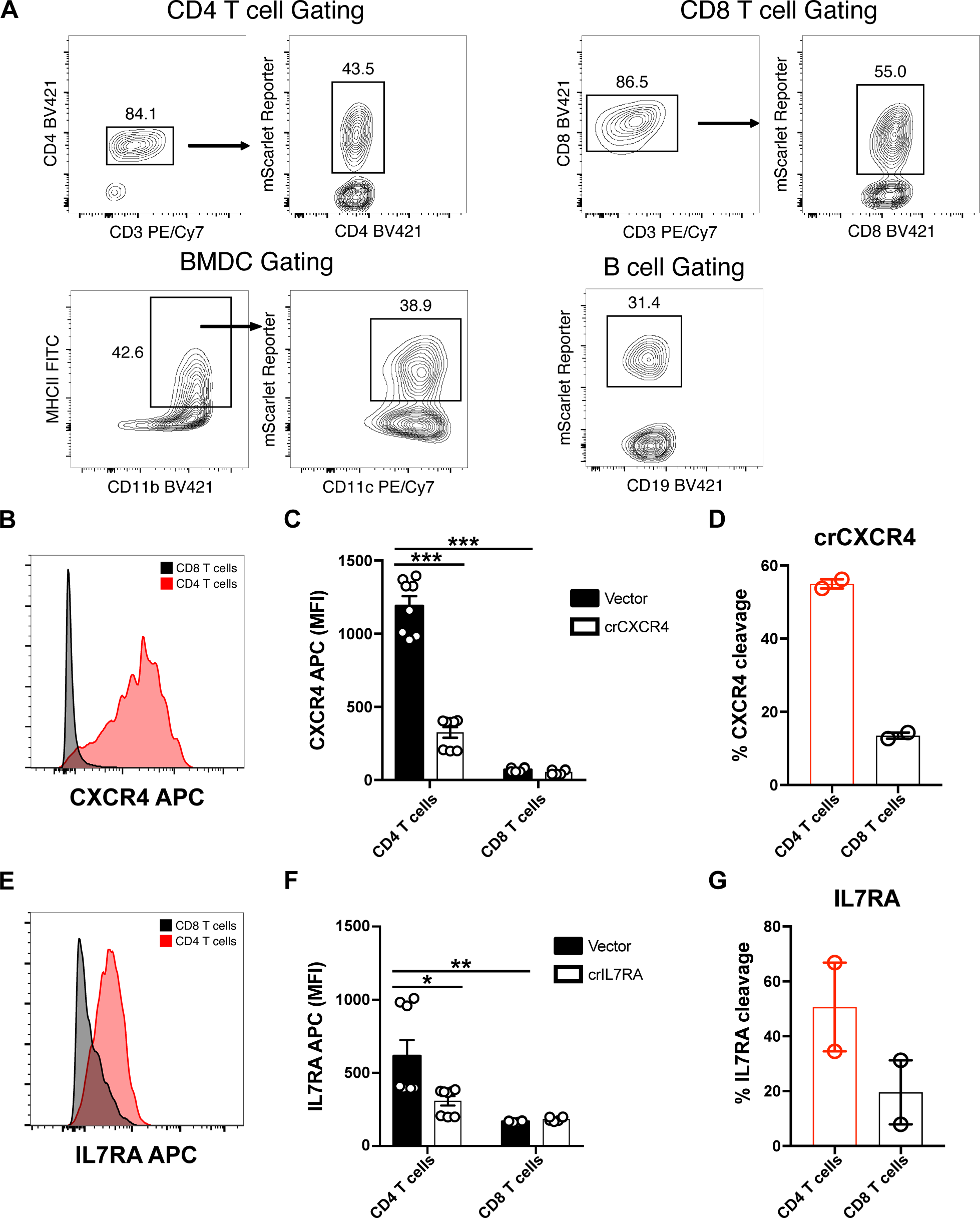
Gating and quantification strategy for charactering gene editing efficiency of primary immune cell. (A) Representative gating schematic used to identify retrovirally infected CD4 T cells (CD3+; CD4+; mScarlet+), CD8 T cells (CD3+; CD8+; mScarlet+), and BMDCs (MCHII+; CD11b+; CD11c+; mScarlet+). (B) Histogram of Cxcr4 expression in CD4 T cells (red) and CD8 T cells (black). (C) Quantification of MFI of Cxcr4 expression in CD4 T cells and CD8 T cells that were either transduced with indicated crRNA or vector control. (D) Quantification of cutting efficiency of each tested crRNA in primary CD4 and CD8 T cells targeting Cxcr4. Data was generated using T7E1 endonuclease surveyor assay. For dot-bar plots, data are shown as mean ± s.e.m. * = p < 0.05, ** = p < 0.01, *** = p < 0.001, by unpaired two-sided t-test. (E) Histogram of Il7ra expression in CD4 T cells (red) and CD8 T cells (black). (F) Quantification of MFI of Il7ra expression in CD4 T cells and CD8 T cells that were either transduced with indicated crRNA or vector control. (G) Quantification of cutting efficiency of each tested crRNA in primary CD4 and CD8 T cells targeting *Il7ra*. Data was generated using T7E1 endonuclease surveyor assay. For dot-bar plots, data are shown as mean ± s.e.m. * = p < 0.05, ** = p < 0.01, *** = p < 0.001, by unpaired two-sided t-test.

**Figure S3.**
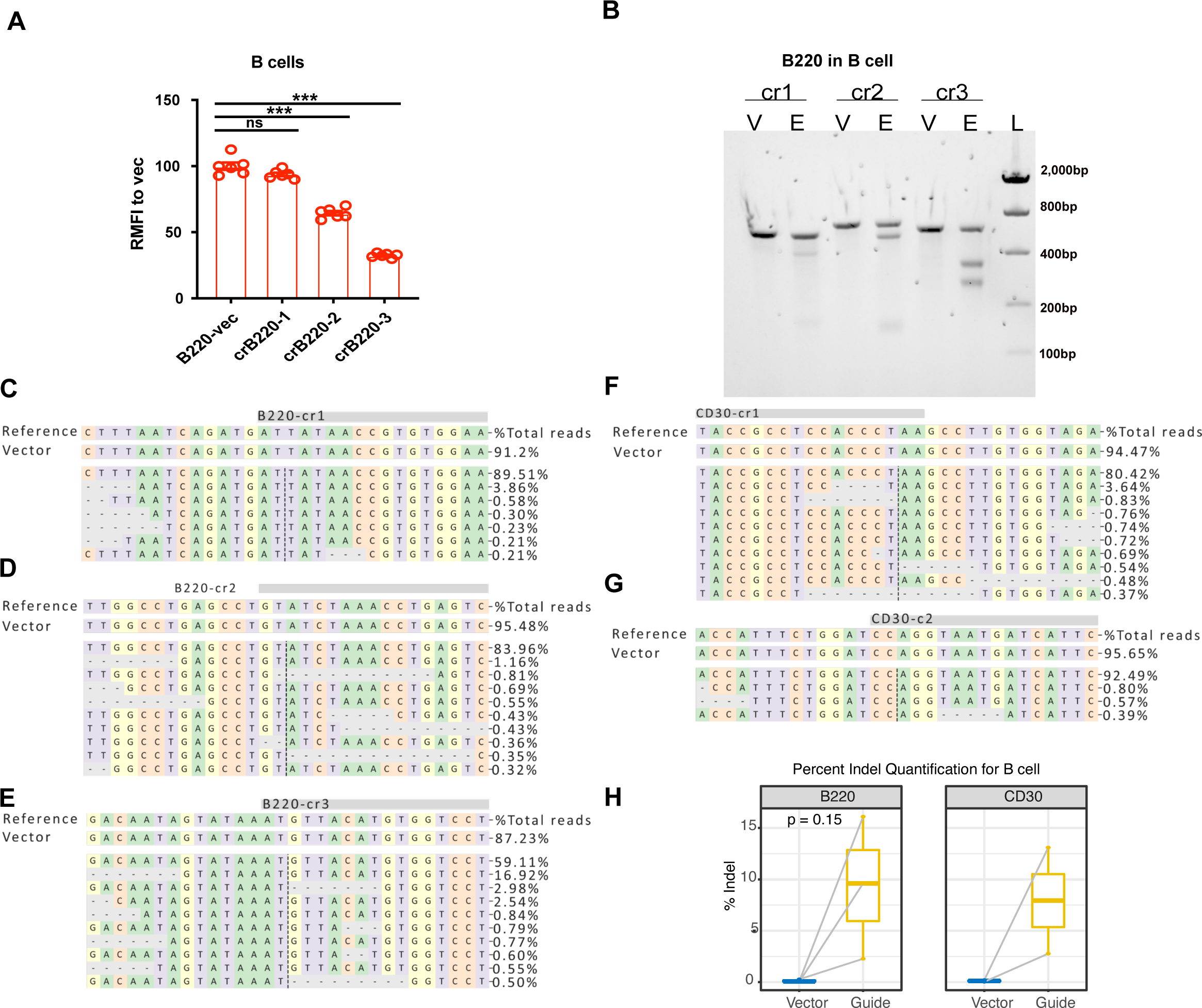
Analysis of gene perturbation in B cells with LbCas12a mice. (A) Quantification of relative mean fluorescence intensity (RMFI) of B220 expression in LbCas12a B cells with respect to the average MFI of vector control. For bar plots, data are shown as mean ± s.e.m. * = p < 0.05, ** = p < 0.01, *** = p < 0.001, by unpaired two-sided t-test. (B) Representative T7E1 gel image demonstrates B220 cleavage in B cells. (C-E) Allele frequency table from the Nextera sequencing of B220 guides. (F-G) Allele frequency table from the Nextera sequencing of CD30 guides. (H) Percent Indel quantification of each guide targeting B220 and CD30 in B cell, compared with its corresponding vector control (represented by two dots connected by grey line). In the box plots, mean from guide group and vector control group are compared by paired two-sided t-test and p-values are displayed on the plots. Multiple guides (crRNAs) are pooled and plotted together.

**Figure S4.**
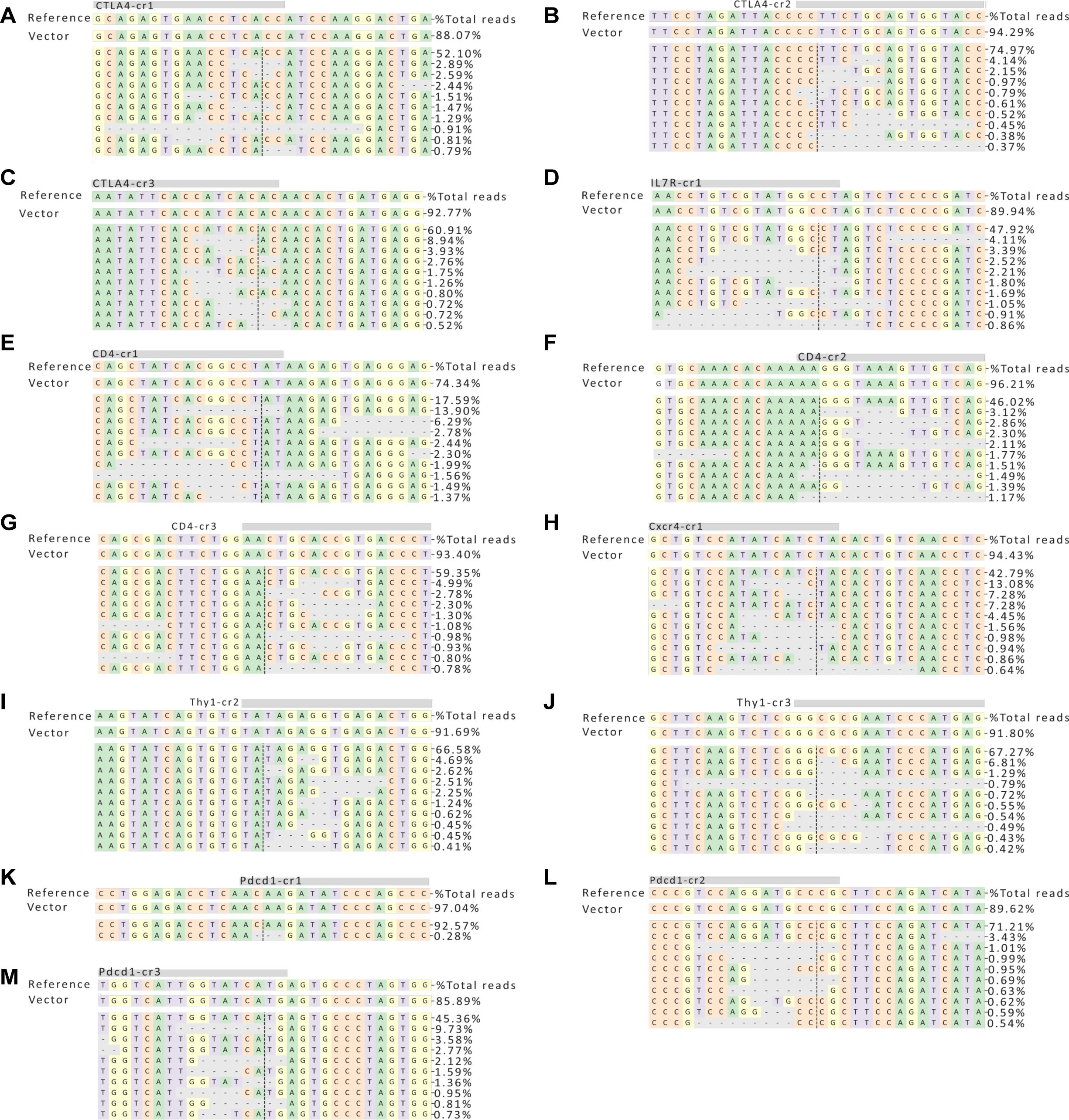
Allele frequency table from Nextera sequencing on CD4 T cells sample. (A-C) Allele frequency table of the *Ctla4* guides (D) Allele frequency table of the *Il7ra* guides (E-G) Allele frequency table of the CD4 guides (H) Allele frequency table of the *Cxcr4* guides (I-J) Allele frequency table of the *Thy1* guides (K-M) Allele frequency table of the *Pdcd1* guides

**Figure S5.**
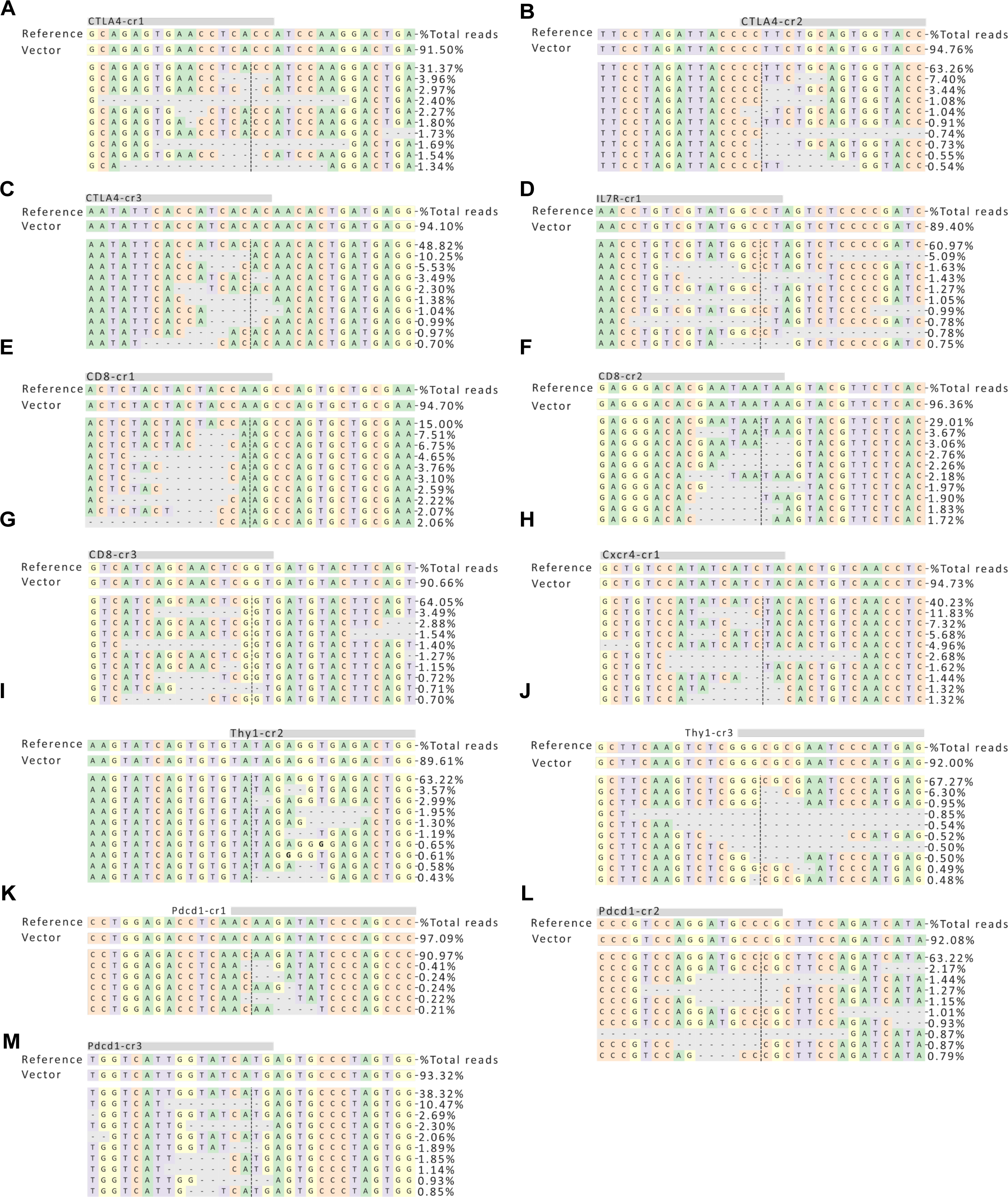
Allele frequency table from Nextera sequencing on CD8 T cells sample. (A-C) Allele frequency table of the *Ctla4* guides (D) Allele frequency table of the *Il7ra* guides (E-G) Allele frequency table of the CD4 guides (H) Allele frequency table of the *Cxcr4* guides (I-J) Allele frequency table of the *Thy1* guides (K-M) Allele frequency table of the *Pdcd1* guides

## Supplemental tables and datasets

**Key Resources Table**

**Table S1. Oligo sequences used in this study**

**Table S2. Aggregated gene editing quantification**

**Dataset S1. Processed NGS data**

## Notes

### Competing Interest Statement

The authors have declared no competing interest.

